# Suppressing STAT3 activity protects the endothelial barrier from VEGF-mediated vascular permeability

**DOI:** 10.1101/2020.10.27.358374

**Authors:** Li Wang, Matteo Astone, Sk. Kayum Alam, Zhu Zhu, Wuhong Pei, David A. Frank, Shawn M. Burgess, Luke H. Hoeppner

**Affiliations:** The Hormel Institute, University of Minnesota, Austin, MN, USA; Translational and Functional Genomics Branch, National Human Genome Research Institute, National Institutes of Health, Bethesda, MD, USA; Department of Medical Oncology, Dana-Farber Cancer Institute, Boston, MA, USA; Masonic Cancer Center, University of Minnesota, Minneapolis, MN, USA

**Author notes:** Corresponding Author: Luke H. Hoeppner, Ph.D., The Hormel Institute, University of Minnesota, 801 16th Avenue NE, Austin, MN 55912. Department of Biology, University of Padova, Padova, Italy (Matteo Astone). These first authors contributed equally (Li Wang, Matteo Astone).

## Abstract

Vascular permeability triggered by inflammation or ischemia promotes edema, exacerbates disease progression, and impairs tissue recovery. Vascular endothelial growth factor (VEGF) is a potent inducer of vascular permeability. VEGF plays an integral role in regulating vascular barrier function physiologically and in pathologies, such as cancer, ischemic stroke, cardiovascular disease, retinal conditions, and COVID-19-associated pulmonary edema and sepsis, which often leads to acute lung injury, including acute respiratory distress syndrome. However, after initially stimulating permeability, VEGF subsequently mediates angiogenesis to repair damaged tissue. Consequently, understanding temporal molecular regulation of VEGF-induced vascular permeability will facilitate developing therapeutics that achieve the delicate balance of inhibiting vascular permeability while preserving tissue repair. Here, we demonstrate that VEGF signals through signal transducer and activator of transcription 3 (STAT3) to promote vascular permeability. Specifically, we show that genetic STAT3 ablation reduces vascular permeability in STAT3-deficient endothelium of mice and VEGF-inducible zebrafish crossed with CRISPR/Cas9 generated genomic STAT3 knockout zebrafish. Importantly, STAT3 deficiency does not impair vascular development and function in vivo. We identify intercellular adhesion molecule 1 (ICAM-1) as a STAT3-dependent transcriptional regulator and show VEGF-dependent STAT3 activation is regulated by JAK2. Pyrimethamine, an FDA-approved anti-microbial agent that inhibits STAT3-dependent transcription, substantially reduces VEGF-induced vascular permeability in zebrafish, mouse, and human endothelium. Indeed, pharmacologically targeting STAT3 increases vascular barrier integrity using two additional compounds, atovaquone and C188-9. Collectively, our findings suggest that the VEGF, VEGFR-2, JAK2, and STAT3 signaling cascade regulates vascular barrier integrity, and inhibition of STAT3-dependent activity reduces VEGF-induced vascular permeability in vertebrate models.

**Key Points:**

- Genetic STAT3 deficiency in VEGF-inducible zebrafish and mice reveals that VEGF signals through STAT3 to promote vascular permeability
- Pyrimethamine, a clinically available agent that inhibits STAT3 activity, reduces VEGF-induced vascular permeability in preclinical models

## INTRODUCTION

Proper physiological function relies on the vascular system to distribute oxygenating blood to all tissues, return deoxygenated blood to the lungs, and maintain tissue homeostasis, including functions such as hemostasis, lipid transport, and immune surveillance. In pathological conditions, vasculature is often adversely affected by the disease process, resulting in vascular permeability^1^. Vascular endothelial growth factor (VEGF) is a central mediator of vascular permeability. In fact, VEGF was initially discovered as a tumor secreted factor that strongly promotes microvascular permeability named ‘vascular permeability factor’^2^ before its subsequent identification as VEGF^3^, an endothelial mitogen essential for the development of blood vessels^4-6^. In cancer, VEGF-induced vascular permeability of plasma proteins creates a matrix amenable to vascular sprouting and tumor growth^2^. In addition to driving tumor angiogenesis, VEGF stimulates tumor cell extravasation, an important step in metastasis that enables cancer cells to enter the bloodstream and potentially invade other tissues^7^. Increased VEGF expression promotes hyperpermeability, edema, and tissue damage leading to the pathogenesis of cardiovascular disease, cerebrovascular conditions, retinal disorders, and acute lung injury. Acute lung injury, including acute respiratory distress syndrome, is among the most severe pathologies caused by coronavirus disease 2019 (COVID-19) and results in pulmonary edema caused by impaired vascular barrier function^8,9^. Autopsy reports of deceased COVID-19 patients commonly describe severe pulmonary mucus exudation, and acute lung injury is frequently a cause of death in severe cases of COVID-19^10,11^. A definitive treatment for acute lung injury does not exist. While therapeutically inhibiting vascular permeability reduces subsequent edema and tissue damage, VEGF-mediated angiogenesis is a key tissue repair mechanism^12,13^. Therefore, temporal VEGF modulation must be achieved when administering therapies to reduce edema and repair ischemic tissue damaged by pathogenesis, which underscores the importance of fully understanding the molecular and temporal regulation of vascular permeability in vivo.

Signal transducer and activator of transcription (STAT) proteins regulate a wide array of cellular functions, including proliferation, differentiation, inflammation, angiogenesis, and apoptosis^14^. Like most of its six other STAT protein family members, STAT3 was identified as part of a cytokine signaling cascade that potentiates the interleukin-6 (IL-6)-mediated hepatic acute phase response as a transcription factor^15,16^. In addition to its prominent role in IL-6 signal transduction, it is well established that VEGF signals through primarily VEGF receptor 2 (VEGFR-2) to stimulate STAT3 activation, dimerization, nuclear translocation, and DNA binding to regulate the transcription of genes involved in endothelial activation, vascular inflammation, and a variety of other biological processes^17-19^. Activation of STAT3 occurs through phosphorylation of tyrosine residue Y705^20^. Many STAT family members are phosphorylated by Janus kinases (JAKs), which are activated through trans-phosphorylation following ligand-mediated receptor multimerization. Mammalian JAK family members include JAK1, JAK2, JAK3 and TYK2^21^. Reports across a variety of tumor and endothelial cell types have suggested members of the Janus kinase (JAK) family, Src, and the intrinsic kinase activity of VEGFR-2 as VEGF-induced activators of STAT3^22-26^. However, the precise mechanism through which VEGF/VEGFR-2 signaling promotes phosphorylation of STAT3 is poorly understood and likely tissue- and cell-type specific.

Here, we identify STAT3 as a central mediator of VEGF-induced vascular permeability. In our study, we exploit the strengths of three model systems, zebrafish, mice, and cultured human endothelial cells, to investigate the role of STAT3 in vascular permeability mediated by VEGF and VEGFR-2 signaling. Among other reasons, zebrafish (*Danio rerio*) have emerged as an invaluable vertebrate model of human pathophysiology due to their genetic similarity to *Homo sapiens* and the transparency of embryos that makes zebrafish amenable to *in vivo* fluorescent imaging^27^. We crossed previously described transgenic heat-inducible VEGF zebrafish^28,29^ to CRISPR/Cas9-generated STAT3 genomic knockout zebrafish^30^ to evaluate the role of STAT3 in VEGF-induced vascular permeability *in vivo*. We establish a complementary VEGF-mediated vascular permeability model in endothelial cell-specific STAT3 knockout mice, demonstrate multiple pharmacological inhibitors of STAT3 reduce vascular permeability *in vivo*, and describe the molecular regulation of VEGF-induced vascular barrier integrity in human endothelial cells.

## METHODS

### Human endothelial cells

Human umbilical vein endothelial cells (HUVEC), human pulmonary artery endothelial cells (HPAEC), and human lung microvascular endothelial cells (HMVEC-L) were certified prior to purchase from Lonza, used exclusively at low passages, and authenticated by morphological inspection.

### Immunoblotting

Human endothelial cells or 3 days post-fertilization (dpf) zebrafish embryos with the yolk sac removed were lysed in RIPA buffer (Millipore) containing protease inhibitors (Roche) and phosphatase inhibitors (Sigma). Proteins were separated via 4-20% gradient SDS-PAGE (Bio-Rad), transferred to membranes, blocked with 5% bovine serum albumin (Sigma-Aldrich), and incubated with primary and secondary antibodies^31^. Antibody-reactive protein bands were detected by enzyme-linked chemiluminescence (Thermo Fisher) using an ImageQuant™ LAS 4000 instrument (GE Healthcare) and quantified using ImageJ software.

### Immunofluorescence

HUVEC, HPAEC, or HMVEC-L were seeded at 4 × 10^4^ cells per well onto EMD Millipore Millicell EZ Slides (PEZGS0416; Sigma Millipore), grown in complete medium for 48 hours, and subsequently serum starved for 16 hours. After receiving inhibitors and/or human recombinant VEGF-165 stimulation, cells were fixed in 4% paraformaldehyde (Boston Bioproducts), permeabilized in cold methanol (Fisher), and immunofluorescence staining was performed. Nuclei were stained using 4’, 6-diamidino-2-phenylindole (DAPI; Cell Signaling; Catalog No. 8961S). All images were captured on a Zeiss Apotome 2 microscope by using 20x, 0.8 NA Plan Apochromat objective magnifying 2X to 4X and processed using ImageJ software (Version 1.8.0_112; https://imagej.nih.gov/ij).

### Vascular permeability assay in mice

Mice with endothelium-specific knockout of STAT3 were created by breeding transgenic mice STAT3^flox/flox^ (Stat3^tm1Xyfu^/J: The Jackson Laboratory) with Tg(Tek-cre)1Ywa/J mice (The Jackson Laboratory). C57BL/6 wildtype mice (Charles River, Catalog No. 027) were purchased and bred. Eight- to ten-week-old pathogen-free male and female mice housed in temperature-controlled room with alternating 12-hour light/dark cycles and fed a standard diet were used for experiments. To assess vascular permeability, Evans Blue dye (100 µl; 1% in PBS; VWR) was intravenously injected in the lateral tail vein of mice. After 15 minutes, mice were anesthetized with ketamine (90-120 mg/kg)/xylazine (5-10 mg/kg) via intraperitoneal injection, and human recombinant VEGF-165 protein (2.5 µg/ml in PBS; MNPHARM; 20 µl total volume; left footpads) and PBS vehicle control (20 µl; right footpads) were each injected into one anterior and one posterior footpad. After 30 minutes, mice were euthanized and footpads were excised. Dye was extracted by incubation in formamide at 63°C overnight and quantified by spectroscopic detection at 620 nm using a Synergy Neo2 instrument (BioTek). Mouse studies were approved by the University of Minnesota (UMN) Institutional Animal Care and Use Committee (IACUC).

### Assessment of vascular permeability in VEGF-inducible, STAT3 deficient zebrafish

Zebrafish were maintained in 28.5°C water and studies were approved by UMN IACUC. Transgenic VEGF-inducible zebrafish^28^ were outcrossed to zebrafish heterozygous for STAT3 deficiency (STAT3^+/-^) generated by CRISPR/Cas9^30^. Subsequently, VEGF-inducible; STAT3^+/-^ zebrafish were incrossed and the 1-cell stage embryos were microinjected with 1.5 nl of Cre mRNA (12.5 ng/µl). Zebrafish expressing the VEGF-inducible transgene were identified by the presence eGFP in their eyes using fluorescent imaging. At 2 dpf, 37°C heat shock of eGFP+ eyed zebrafish carrying the VEGF transgene was performed to confirm VEGF transgene activity via the absence of mCherry fluorescence. At 3 dpf, zebrafish were anesthetized and fluorescent microangiography was performed. A microneedle was inserted through the pericardium directly into the ventricle, and a mixture of 2000 kDa FITC-dextran and 70 kDa Texas Red-dextran (2 mg/ml in embryo water; Life Technologies, Inc.) was injected. Immediately prior to imaging, 37°C heat shock induction of the VEGF transgene was performed for 10 minutes. Real-time imaging using SCORE methodology^32^ was performed using a Zeiss Apotome 2 microscope.

### In vitro kinase assay

Human JAK2 protein with active kinase activity (Signal Chem; J02-11G-05) and purified human STAT3 protein from Sf9 cells were subjected to an in vitro kinase assay using previously described methods^33^. Briefly, 10 µl JAK2 diluted in kinase dilution buffer III (Signal Chem; K23-09) to a final concentration of 0.1µg/ml was incubated with 3 µg purified STAT3 proteins as well as 5 µl ATP (New England Biolabs; N0440S) for 30 minutes at 30°C. The incubation was terminated by boiling the samples at 95^°^C for 5 minutes in 1X Laemmli sample buffer (Bio-Rad) supplemented with 10% β-mercaptoethanol (Bio-Rad). The samples were analyzed by immunoblotting using anti-phosphorylated STAT3 antibody (Y705; Cell signaling Technology) to validate JAK2-mediated kinase activity upon STAT3 protein at the Tyr705 position.

### Dual-luciferase reporter assay

pGL3-ICAM1 luciferase reporter (pGL3-ICAM1-WT) was a generous gift from Dr. Jim Hu at the Hospital for Sick Children in Toronto, Ontario, Canada^34^. QuikChange Lightning Site-Directed Mutagenesis (Aligent) was performed as described^35^ to mutate the STAT3 binding site within the ICAM1 promoter from 5’-TTC-CxG-GAA-3’ to 5’-AGC-CxC-CTG-3’. Constitutively active STAT3 plasmid EF.STAT3C.Ubc.GFP was a gift from Dr. Linzhao Cheng (Addgene plasmid # 24983; http://n2t.net/addgene:24983; RRID: Addgene_24983)^36^. HUVEC seeded at 1.5 × 10^6^ cells per well of a collagen-coated 6-well pate were transfected with plasmids using the Neon Transfection System (Invitrogen), harvested 48 hours post-transfection, Dual-Luciferase Reporter Assays (Promega) were performed, and firefly and renilla luciferase luminescence was measured using a Synergy Neo 2 Reader (BioTek).

### Statistical analysis

Unpaired Student’s t-test, paired t-test, or ANOVA was used to compare differences between groups as indicated and values of P < 0.05 were considered significant. Data are expressed as mean ± SEM and representative of at least three independent experiments.

### Data sharing statement

In addition to data reported in the manuscript and Supplemental Figures, any datasets used and/or analyzed during the current study are available from the corresponding author upon request.

## RESULTS

### VEGF/VEGFR-2 induces STAT3 phosphorylation and nuclear localization

Given that we sought to investigate STAT3 as an important regulator of VEGF-induced vascular permeability, we first performed studies to confirm VEGF activates STAT3 in endothelium. We observed activation of VEGFR-2 and STAT3 following VEGF stimulation (**Figure 1A**), which coincides with prior reports^23^. VEGF stimulation promotes physical interaction of VEGFR-2 and STAT3, as evident by immunoprecipitation studies demonstrating an association of total VEGFR-2 and total STAT3 (**Figure 1B**) as well as VEGF-dependent interactions between the phosphorylated forms of VEGFR-2 and STAT3 (**Figure 1C**). To further study the interaction between VEGFR-2 and STAT3, we performed a GST pull-down experiment using GST-tagged STAT3 protein as a “bait” protein and growth factor stimulated HUVEC lysate as a source of “prey” proteins. We observed a growth factor-dependent interaction between endothelial cell-derived VEGFR-2 and GST-tagged STAT3 proteins (**Figure 1D**). Immunofluorescence studies demonstrate that STAT3 translocates to the nucleus upon VEGF/VEGFR-2-mediated activation in HUVEC (**Figure 1E-F**). Collectively, our results suggest VEGF stimulates VEGFR-2 to induce STAT3 phosphorylation and nuclear localization in human endothelial cells.

**Figure 1:**
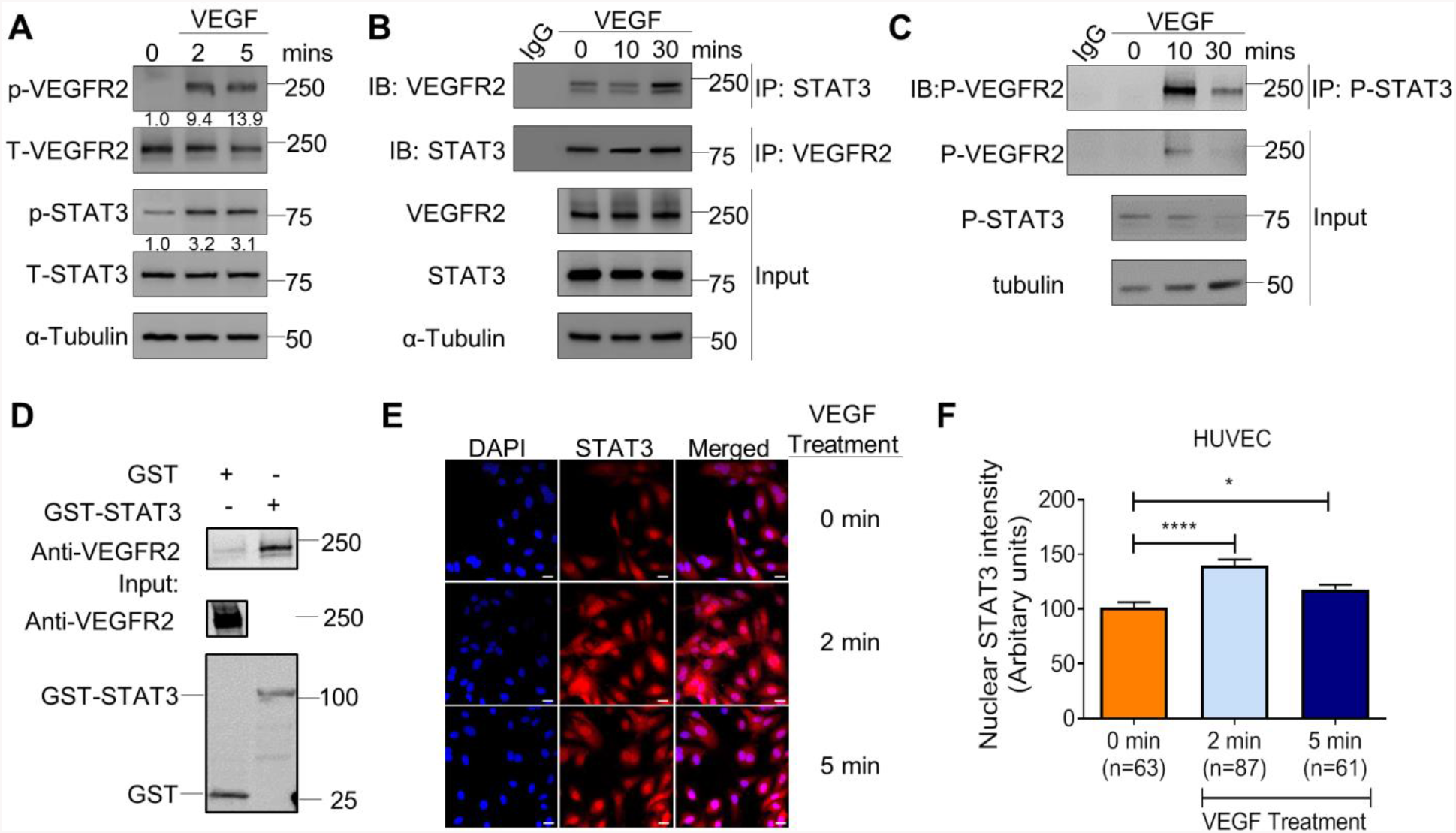
VEGF/VEGFR-2 induces STAT3 phosphorylation and nuclear localization. (A) Stimulation of HUVEC with 25ng/ml recombinant human VEGF-165 protein for 2 and 5 minutes induces p-VEGFR-2 (Y1175) and p-STAT3 (Y705) via immunoblotting. (B) VEGF (25 ng/ml) stimulation in HUVEC for 10 and 30 minutes promotes co-immunoprecipitation of STAT3 and VEGFR-2. (C) VEGF (25 ng/ml) stimulation in HUVEC for 10 and 30 minutes promotes co-immunoprecipitation of p-STAT3 (Y705) and p-VEGFR-2 (Y1175). (D) GST pull-down of VEGFR-2 with STAT3. Lysates of HUVEC stimulated with serum for 30 minutes were used as prey. GST fusion protein STAT3 expressed in 293F cells was used as bait. GST alone served as a negative control. Binding experiments were analyzed by SDS-PAGE and visualized by immunoblot. GST-STAT3 and GST were both detected using an anti-GST antibody. (E) VEGF stimulation for 2 minutes and 5 minutes promotes nuclear localization of STAT3. DAPI is blue. Scale bar, 20 µm. (F) Quantification of nuclear immunofluorescence staining intensity. Mean ±SEM, one-way ANOVA. *P<0.05, ****<0.0001. (A-E) Images are representative of multiple biological replicates.

### VEGF-induced vascular permeability is reduced in STAT3 deficient zebrafish

To investigate the role of STAT3 in VEGF-induced vascular permeability *in vivo*, we utilized a heat shock inducible VEGF transgenic zebrafish (iVEGF) that we previously developed to identify regulators of VEGF-mediated vascular leakage^28,29^. Specifically, we generated STAT3 knockout zebrafish with an inducible VEGF transgene by crossing iVEGF zebrafish^28^ to CRISPR/Cas9-generated STAT3 genomic knockout (STAT3^KO^) zebrafish (**Figure 2A**). Importantly, we did not observe any vascular development defects in STAT3^KO^ larval zebrafish, and the vascular system of STAT3^KO^ zebrafish at 3 dpf is indistinguishable from wildtype STAT3^+/+^ siblings (**Figure 2B**). To assess vascular permeability, VEGF was heat-induced in STAT3^KO^; iVEGF zebrafish following ventricular co-injection of 70 KDa Texas Red-dextran as a permeabilizing tracer and 2000 KDa FITC-dextran as a marker of the veins. Zebrafish were immediately live imaged to measure vascular permeability, evident by leakage of Texas Red-dextran into the extravascular space^28^. Using these techniques, our data showed decreased VEGF-induced vascular permeability in STAT3^KO^ zebrafish relative to corresponding wildtype controls (**Figure 2C-D**), suggesting that VEGF signals through STAT3 to promote vascular permeability.

**Figure 2:**
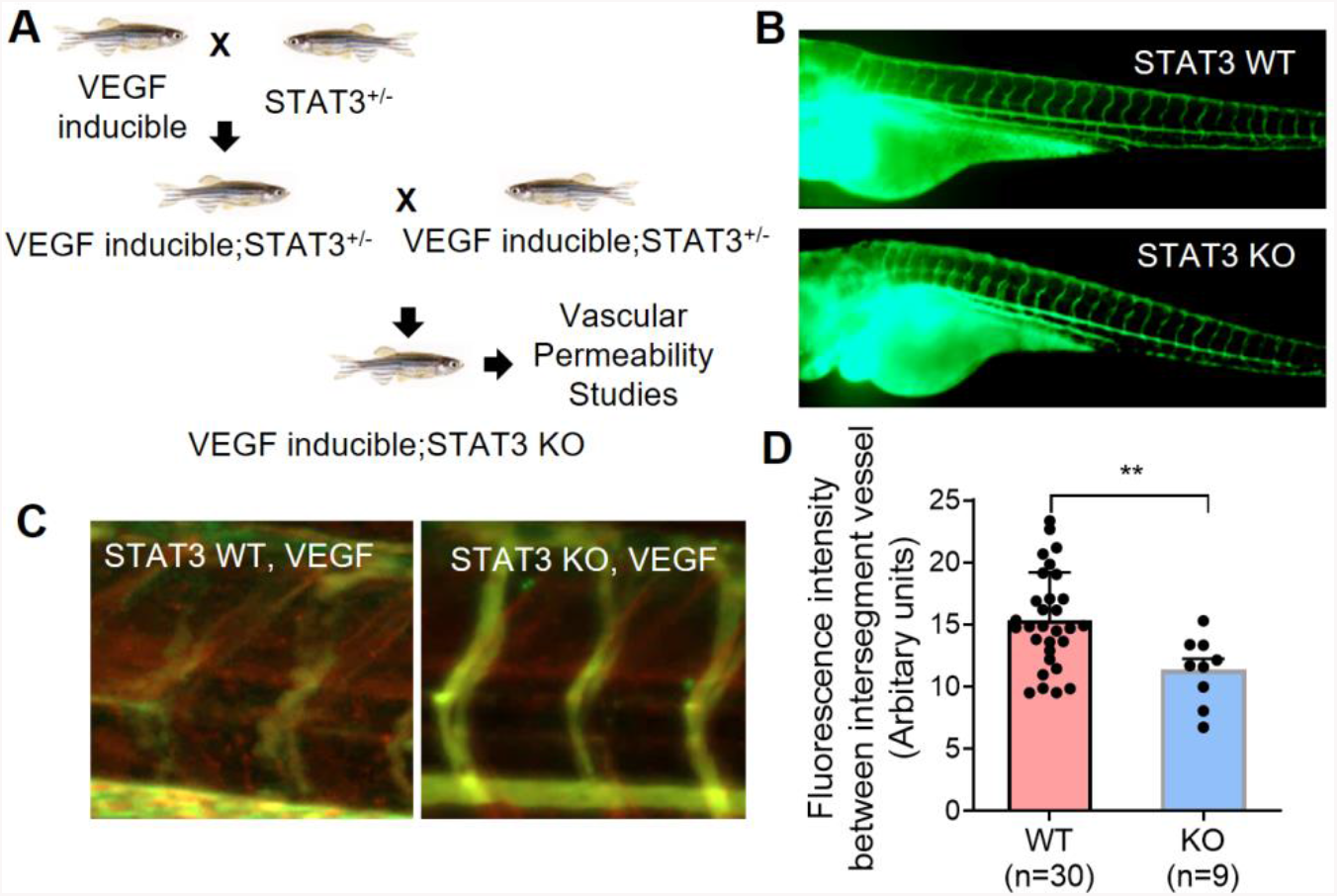
VEGF-induced vascular permeability is reduced upon CRISPR-mediated knockout of STAT3 in zebrafish. (A) VEGF-inducible zebrafish were crossed to STAT3+/- (heterozygous) zebrafish to generate VEGF-inducible; STAT3+/- double transgenic fish, which were intercrossed to generate VEGF-inducible; STAT3-/- (KO) zebrafish. (B) CRISPR/Cas9-generated STAT3 KO zebrafish display no overt vascular defects relative to WT. Vascular system visualized by microangiography with 2000 kDa FITC-dextran. (C) Microangiography using 70 kDa Texas Red-dextran permeabilizing tracer (red) and 2000 kDa FITC-dextran ISV marker (green) was performed on 3 days old VEGF-induced, STAT3^**+/+**^ (n=30) and VEGF-induced, STAT3^**-/-**^ (n=9) zebrafish. (D) Quantitative analysis of vascular permeability upon VEGF stimulation in wildtype (STAT3^**+/+**^) and knockout (STAT3^**-/-**^) zebrafish. **P<0.01, unpaired t-test (B, C) Representative images shown were obtained using a Zeiss Apotome 2 microscope with a Fluar 5x, 0.25 NA lens at RT. ISV: inter-segmental vessel.

### Endothelial-specific STAT3 knockout mice exhibit decreased VEGF-induced vascular permeability

To corroborate findings from the zebrafish vascular permeability model in a mammalian system, we have standardized a mouse footpad permeability assay in endothelial cell specific-STAT3 deficient mice (STAT3^ECKO^). Given that germline STAT3 deficiency leads to embryonic lethality in mice^37^, we generated STAT3^ECKO^ mice by crossing Tie2-Cre mice^38^ to STAT3 floxed mice^39^. To assess VEGF-induced vascular permeability, anesthetized STAT3^ECKO^ or corresponding control mice were injected intravenously with Evans blue dye, and subcutaneously injected with recombinant VEGF protein (2.5 µg/ml in PBS; left footpads) or vehicle (right footpads). After 30 minutes, footpads were excised from euthanized mice, extravasated dye was extracted via formamide, and measured by spectrometry to quantify VEGF-induced vascular permeability. We observed significantly decreased extravasation of Evans blue dye in STAT3^ECKO^ mice relative to controls, suggesting STAT3 is an important transducer of VEGF-induced vascular permeability (**Figure 3**).

**Figure 3:**
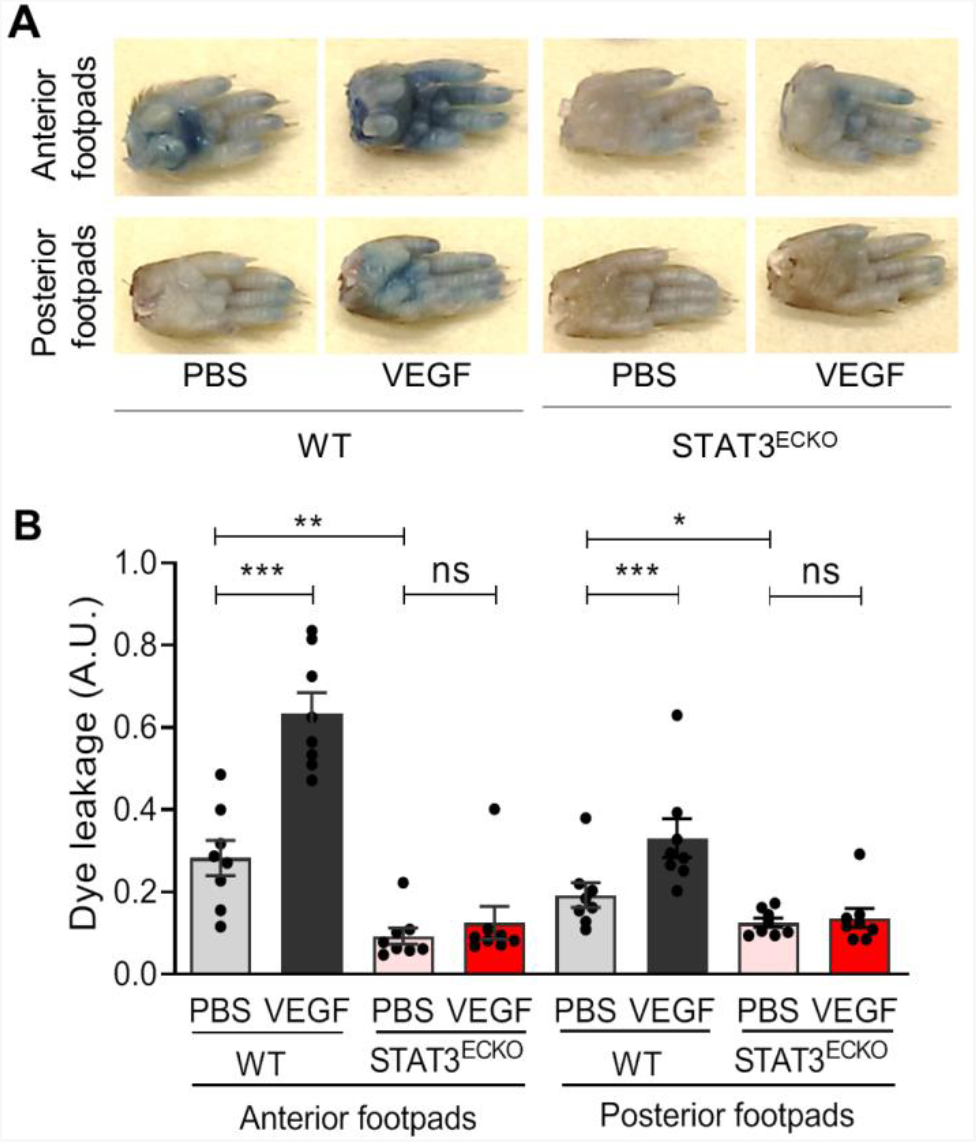
Endothelial cell-specific STAT3 knockout mice exhibit decreased VEGF-induced permeability. (A) Images of footpads from WT and STAT3^ECKO^ (endothelial cell-specific STAT3 KO) mice following tail vein injection with 1% Evans blue and human recombinant VEGF-165 protein (2.5 µg/ml) or PBS vehicle being injected into the root of the footpad. (B) Quantitation of Evans blue leakage in Tie2-Cre negative; STAT3^flox/flox^ (WT) and Tie2-Cre positive; STAT3^flox/flox^ (STAT3^ECKO^) mice. n=8 mice per group. Each mouse was injected with PBS on right anterior and posterior footpads and VEGF on left anterior and posterior footpads. Multiple biological replicates were performed and depicted findings are representative. *P<0.05, **P<0.01, ***P<0.001, one-way ANOVA followed by Dunnett’s test.

### Pharmacological inhibition of STAT3 stabilizes endothelial barrier integrity following VEGF stimulation

Given that genetic ablation of STAT3 reduces vascular permeability in zebrafish and mice, we sought to test the effects of pharmacological STAT3 inhibition on endothelial barrier integrity. Pyrimethamine (PYR; Daraprim®)^40^ and Atovaquone (AQ; Mepron®)^41^ are FDA-approved antimicrobial agents that were recently discovered as new inhibitors of STAT3 activity. PYR was identified as a STAT3 inhibitor through a chemical biology approach. AQ rapidly and specifically downregulates cell-surface expression of glycoprotein 130, which is required for STAT3 activation in multiple contexts. These compounds have been shown to be safe in humans because they inhibit STAT3 at concentrations routinely achieved in human plasma^40-42^. To our knowledge, PYR and AQ have yet to be specifically assessed in endothelium as prior studies have primarily evaluated their STAT3 inhibitory effects on tumor cells of epithelial origin^40,41,43-49^. Therefore, we first treated HUVEC with various concentrations of PYR or AQ for different durations of time to determine the optimal conditions, while not exceeding the concentrations typically achieved in human plasma **(Supplemental Figure 1)**. In serum-starved HUVEC treated with 10 µM PYR for 1 hour or 30 µM AQ for 4 hours followed by VEGF stimulation, we observed decreased phosphorylation of STAT3 at Y705 **(Figure 4A-B, Supplemental Figure 2)**. We confirmed activation of VEGF signaling by assessing the phosphorylation status and total protein levels of VEGFR-2 and JAK family members **(Figure 4B, Supplemental Figure 2)**. Given that VEGFR-2, JAK1, JAK2, and TYK2 are upstream of STAT3, PYR- and AQ-mediated STAT3 inhibition does not affect activation of these proteins. Taken together, these results suggest PYR and AQ inhibit VEGF-induced STAT3 activation in human endothelial cells.

**Figure 4:**
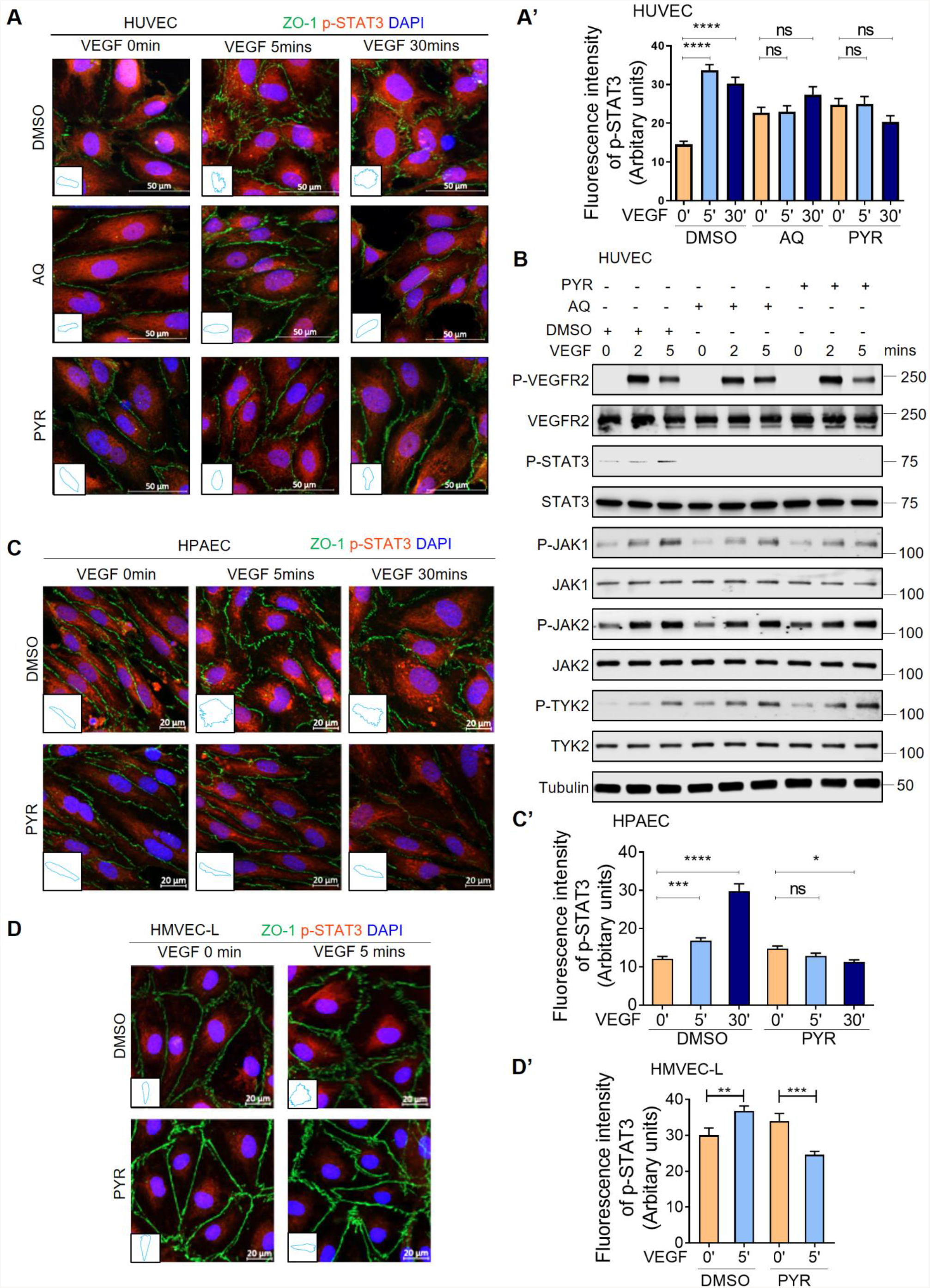
Pharmacological inhibition of STAT3 stabilizes endothelial barrier integrity following VEGF stimulation in human endothelial cells. (A) Human VEGF-165 recombinant protein (VEGF; 25 ng/ml) stimulation of HUVEC promotes ZO-1 (green) disorganization at endothelial cell junctions (top: DMSO vehicle control pretreatment for 1 hour prior to VEGF stimulation). ZO-1 organization is maintained upon pretreatment with 30 μM AQ for 4 hours (middle) or 10 μM PYR for 1 hour (bottom) prior to VEGF stimulation. VEGF-induced phosphorylation of STAT3 at Y705 (p-STAT3; red) was reduced upon AQ or PYR pretreatment. Nuclei: DAPI (blue). Insets: ZO-1 staining trace of 1 representative cell/field. (A’) Quantification of p-STAT3 (red). (B) Serum-starved HUVEC were pretreated with DMSO (vehicle control) for 1 hour, 30 µM AQ for 4 hours, or 10 µM PYR for 1 hour prior to VEGF (25 ng/ml) stimulation for 0, 2 or 5 minutes. Lysates were immunoblotted. Please see Supplemental Figure 2 for densitometry analysis. (C) Serum-starved HPAEC were pretreated with 10 µM PYR for 1 hour prior to VEGF (25 ng/ml) stimulation for 0, 5 or 30 minutes. VEGF stimulation promotes disorganization of ZO-1 (green) at endothelial cell junctions. ZO-1 organization is maintained when HPAEC were pretreated with PYR. VEGF-induced p-STAT3 (red) was reduced upon PYR pretreatment. Nuclei were stained with DAPI (blue). Insets: trace of ZO-1 staining on 1 representative cell per field. (C’) Quantification of p-STAT3 (red). (D) VEGF (25 ng/ml) stimulation of HMVEC-L promotes ZO-1 (green) disorganization at endothelial cell junctions. ZO-1 organization is maintained upon pretreatment with 20 μM PYR for 6 hours prior to VEGF stimulation. VEGF-induced p-STAT3 (red) was reduced upon PYR pretreatment. Nuclei: DAPI (blue). Insets: ZO-1 staining trace of 1 representative cell/field. (D’) Quantification of p-STAT3 (red). ****P<0.0001, ***P<0.001, **P<0.01, *P<0.05, one-way ANOVA.

As vascular barrier stability is in large part regulated by intercellular junctions^50^, we examined whether STAT3 inhibition affects junctional organization in human endothelial cells. In HUVEC, HPAEC, and HMVEC-L, we demonstrate by immunofluorescence that VEGF induces STAT3 activation via Y705 phosphorylation and zonula occludens 1 (ZO-1) disorganization, indicative of tight junction instability (**Figure 4A, C, D**). STAT3 pharmacological inhibition via PYR or AQ restores ZO-1 organization in HUVEC, suggesting that VEGF may mediate vascular permeability through STAT3-regulated control of ZO-1 (**Figure 4A**). The frequency of tight junctions observed by transmission electron microscopy has been shown to increase twofold by culturing HUVEC in a 1:1 mixture of astrocyte-conditioned medium and standard endothelial medium^51^. In HUVEC cultured in the presence astrocyte-conditioned medium, we observe VEGF-mediated activation of VEGFR-2, JAK2, and STAT3 with increased ZO-1 disorganization, while PYR and AQ inhibit STAT3 phosphorylation and restore tight junction stability through proper ZO-1 organization (**Supplemental Figures 3 & 4**), which coincides with our findings from human endothelial cells cultured in standard medium. PYR pretreatment also prevented VEGF-induced ZO-1 disorganization in HPAEC and HMVEC-L cultured in standard conditions (**Figure 4C-D**). Correspondingly, we demonstrate that C188-9, a high-affinity inhibitor of STAT3 that targets its phosphotyrosyl peptide binding site within the SRC homology 2 (SH2) domain^52^, prevents VEGF-mediated ZO-1 disorganization in HUVEC and HPAEC (**Supplemental Figure 5A-D**).

Given that genetic ablation of STAT3 in zebrafish and mice reduces VEGF-mediated vascular permeability, we next sought to assess the effects of pharmacological STAT3 inhibition on endothelial barrier integrity *in vivo*. To that end, VEGF-inducible zebrafish embryos were exposed to 10 µM PYR via their water for 3 days and vascular permeability was assessed using fluorescent microangiography. We observed decreased VEGF-induced vascular permeability in zebrafish treated with PYR relative to controls, suggesting that pharmacological inhibition of STAT3 reduces VEGF-induced vascular permeability in zebrafish (**Figure 5 A-B; Supplemental Figure 6**). To evaluate permeability upon STAT3 inhibition in mice, we treated wildtype C57BL/6 mice with 75 mg/kg PYR daily for 15 consecutive days, 100 mg/kg C188-9 daily for 7 consecutive days, or each corresponding vehicle control via intraperitoneal injection, followed by vascular permeability assessment as described for similar experiments. We observed decreased VEGF-induced extravasation of dye in mice treated with STAT3 inhibitor relative to controls, suggesting that pharmacological inhibition of STAT3 reduces vascular permeability in mice **(Figure 5C-D, Supplemental Figure 5E)**.

**Figure 5:**
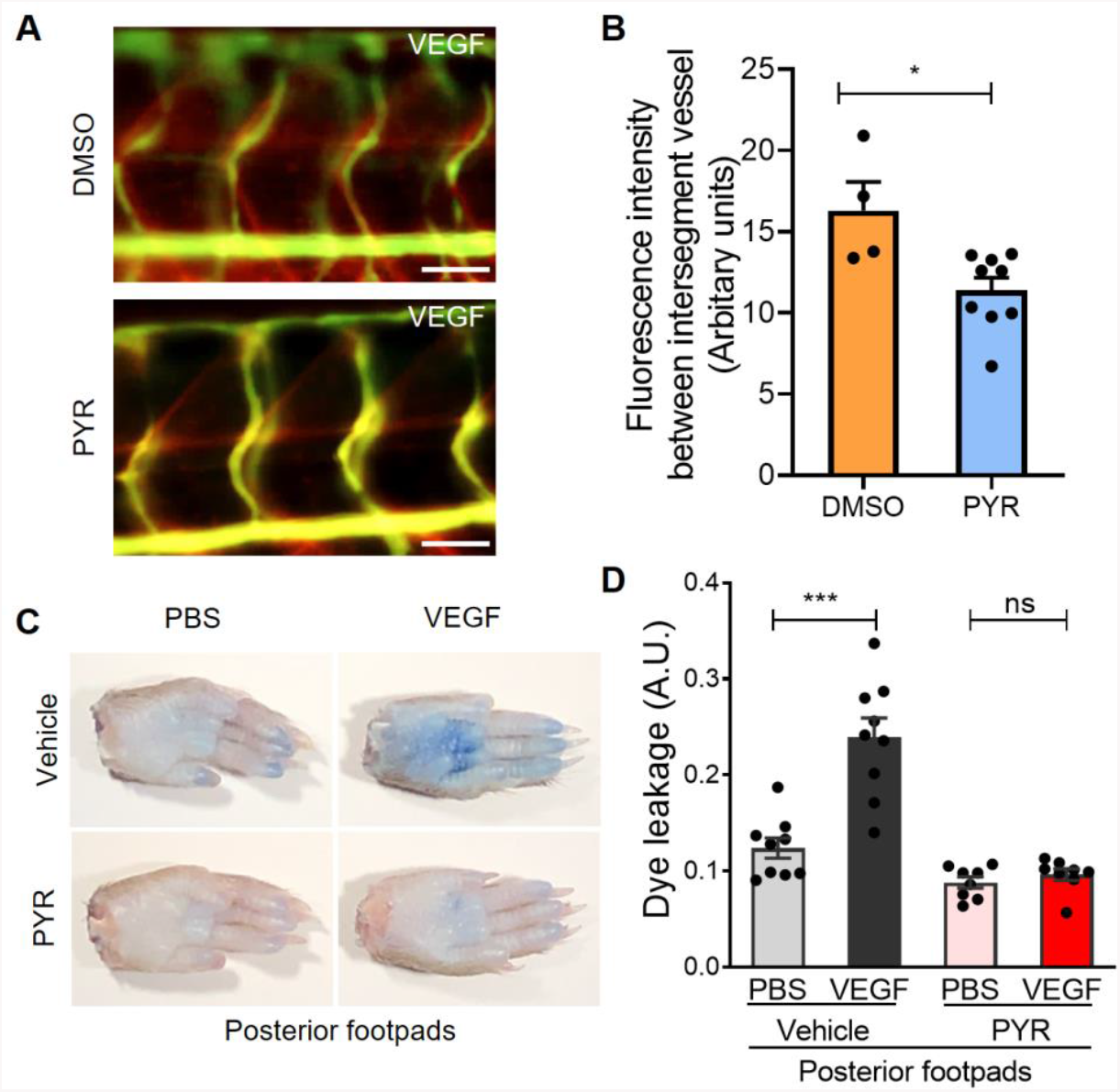
Suppression of STAT3 activity by pyrimethamine inhibits VEGF-induced vascular permeability in zebrafish and mice. (A) Microangiography using 70 kDa Texas Red-dextran permeabilizing tracer (red) and 2000 kDa FITC-dextran ISV marker (green) was performed on 3 day old VEGF-induced zebrafish pretreated with DMSO (n=4) or 25 μM PYR (n=9) for 3 days. Representative images shown were obtained using a Zeiss Apotome 2 microscope with a Fluar 5x, 0.25 NA lens at RT. ISV: intersegmental vessel. Scale bar: 50 μm. (B) The quantitative analysis of vascular permeability upon VEGF stimulation in zebrafish pretreated with DMSO or PYR. *P<0.05, unpaired t-test. (C) Representative images of footpads from mice treated with vehicle or PYR following tail vein injection with 1% Evans blue and footpad injection of VEGF (2.5 μg/ml) or PBS vehicle. (D) Quantitation of Evans blue leakage in C57BL/6 wildtype mice treated with vehicle or PYR. n=9 mice in the vehicle group and n=8 mice in the PYR group. Each mouse was injected with PBS on right anterior and posterior footpads and VEGF on left anterior and posterior footpads. Multiple biological replicates were performed and depicted findings are representative. ***P<0.001, paired t-test.

### JAK2 activates STAT3 to promote VEGF/VEGFR-2-induced vascular permeability

Phosphorylation of STAT3 at Y705 causes its dimerization, nuclear translocation, and DNA binding to control transcription of genes, including regulators of vascular permeability. The kinase(s) that phosphorylates STAT3 Y705 in endothelial cells has not been well-identified. VEGF signals through VEGFR-2, which possesses intrinsic kinase activity through which it may activate STAT3 directly or through another kinase, such as JAK1, JAK2, JAK3, TYK2, or SRC^22-24,26^. Here, our findings suggest that JAK2 phosphorylates STAT3 in HUVEC and show JAK2 is required for VEGF/VEGFR-2-mediated vascular permeability *in vivo*. Specifically, we first tested whether JAK2 physically interacts with STAT3. Indeed, in GST pulldown assays purified GST-STAT3 fusion protein associates with each of VEGFR-2 and JAK2 present in lysates derived from stimulated HUVEC **(Figure 6A)**. To determine whether JAK2 directly phosphorylates STAT3, we performed an in vitro kinase assay using kinase active JAK2 protein with STAT3 protein we purified from Sf9 insect cells and found that JAK2 phosphorylates STAT3 at Y705 **(Figure 6B)**. To test the role of JAK2 in the regulation of vascular barrier integrity, we administered a JAK2 inhibitor, AG490, to C57BL/6 mice daily for seven consecutive days and assessed VEGF-induced vascular permeability of intravenously injected Evans blue dye. Pharmacological inhibition of JAK2 substantially reduced VEGF-mediated vascular permeability in mice **(Figure 6C-D)**, suggesting vascular permeability induced by VEGF/VEGFR-2 requires JAK2. Based on our collective findings, JAK2 activates STAT3 via Y705 phosphorylation to promote VEGF/VEGFR-2-induced vascular permeability.

**Figure 6:**
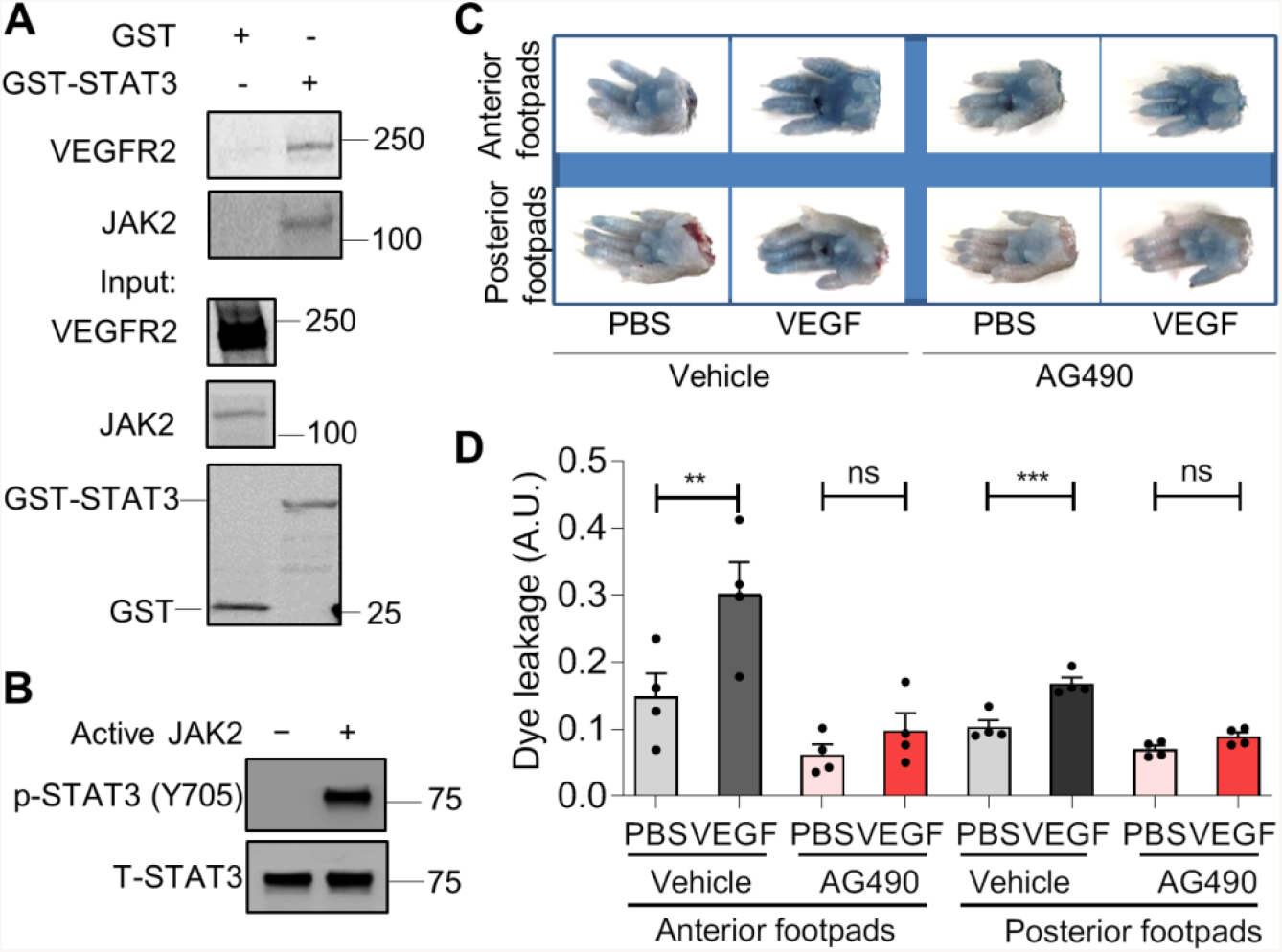
JAK2 phosphorylates STAT3 to transduce VEGF/VEGFR-2 signaling and promote vascular permeability. (A) To perform a STAT3 GST pull-down of VEGFR-2 and JAK2, lysates of HUVEC stimulated with serum for 30 minutes were used as prey. GST fusion protein STAT3 expressed in 293F cells was used as bait. GST alone served as a negative control. Binding experiments were analyzed by SDS-PAGE and visualized by immunoblotting. GST-STAT3 and GST were each detected using an anti-GST antibody. (B) JAK2 phosphorylates STAT3 in vitro. In vitro kinase assay was performed using purified human STAT3 protein and kinase active JAK2 protein. (C) Images of footpads from C57BL/6 wildtype mice treated with vehicle or JAK2 inhibitor AG490. Following tail vein injection with 1% Evans blue, human VEGF-165 protein (2.5 μg/ml) or PBS vehicle were injected into the root of the footpad. After 30 minutes, the mice were euthanized and the footpads were excised. (D) Quantitation of Evans blue leakage in C57BL mice treated with vehicle or AG490. n=4 mice per group. Each mouse was injected with PBS on right anterior and posterior footpads and VEGF on left anterior and posterior footpads. **P<0.01, ***P<0.001, paired t-test.

### STAT3 transcriptionally activates ICAM1, a cell adhesion molecule that promotes vascular permeability

We next sought to investigate molecular mechanisms through which STAT3 transcriptionally regulates VEGF-induced vascular permeability. To that end, we identified a novel STAT3 binding site within the promoter region of ICAM-1 **(Figure 7A)**, a cell adhesion molecule known to promote vascular permeability. Using luciferase-based reporter assays, we demonstrate increased ICAM-1 promoter activity upon co-transfection with increasing amounts of constitutively active STAT3 cDNA plasmid (**Figure 7B**). Mutation of the STAT3 binding site within the ICAM-1 promoter region prevents activation of the ICAM-1 promoter by constitutively active STAT3 (**Figure 7C**). Finally, we confirmed that ICAM-1 protein is upregulated following VEGF-mediated activation of STAT3 in human endothelial cells **(Figure 7D)**. Taken together, our findings suggest VEGF-induced STAT3 transcriptionally regulates ICAM-1 in human endothelium.

**Figure 7:**
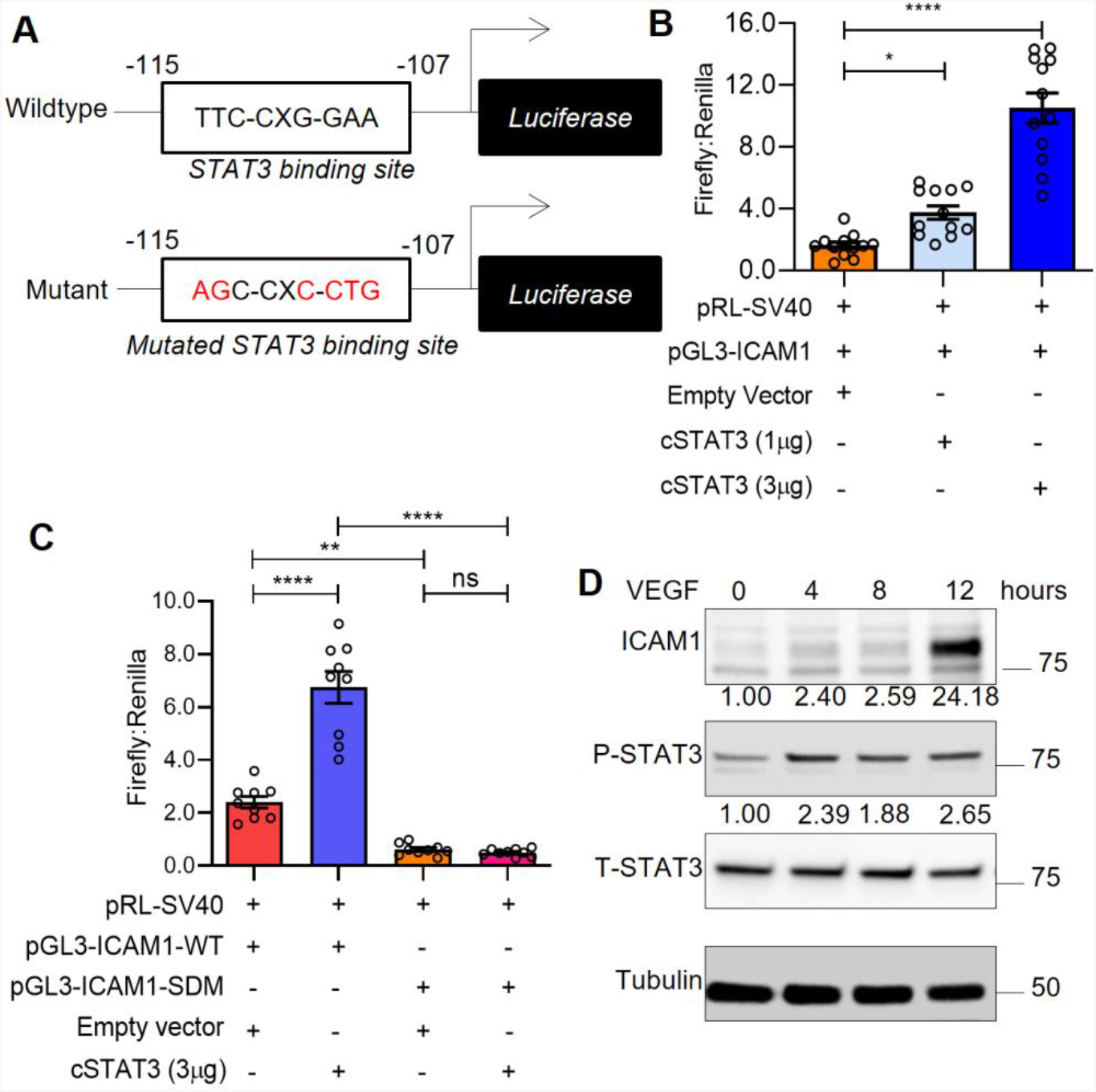
STAT3 transcriptionally activates ICAM1, a cell adhesion molecule that promotes vascular permeability. (A) The pGL3-ICAM1-WT plasmid (top) containing the human ICAM1 promoter with a STAT3 binding site located at −115 to −107bp. The pGL3-ICAM1-SDM plasmid (bottom) with mutation in the STAT3 binding site as indicated. (B) HUVEC were transiently transfected via electroporation with pGL3-ICAM1-WT (Firefly), pRL-SV40 (Renilla) plasmids and different amounts of constitutively active STAT3 plasmid (1 µg and 3 µg) using Neon transfection system. Firefly and Renilla luminescence was measured and plotted as ratio. Mean ±SEM, two-tailed unpaired t-test. n=12 technical replicates. *P<0.05, ****P<0.0001. (C) Dual-luciferase assays were performed in HUVEC that were transfected with pGL3-ICAM1-WT or pGL3-ICAM1-SDM and empty vector or constitutively active STAT3. Firefly and Renilla luminescence was measured and plotted as a ratio. Mean ±SEM, two-tailed unpaired t-test. n=9 technical replicates. **P<0.01, ****P<0.0001. (D) Human VEGF-165 protein (25ng/ml) stimulated HUVEC lysates were immunoblotted for ICAM1, p-STAT3 (Y705), and total STAT3. (B-D) Depicted data is representative of multiple biological replicates. SDM: Site-directed mutagenesis.

## DISCUSSION

Despite the prominent role of VEGF-induced vascular permeability in pathogenesis, a greater collective molecular understanding of its regulatory mechanisms is necessary to therapeutically improve vascular barrier integrity to prevent subsequent edema and tissue damage in vascular diseases like myocardial infarction, ischemic stroke, and acute lung injury. These research efforts have been hindered by limitations and inconsistencies in current accessible models. Endothelial cell culture models often fail to uniformly replicate in vivo models of hyperpermeability^53,54^. Molecular regulation of vascular barrier integrity can vary greatly based on various pathophysiological contexts and differences in cell types, tissues, anatomical locations, and species^55^. To overcome these obstacles, we took advantage of a transgenic VEGF-inducible zebrafish model that is amenable to genetic manipulation and reproducible live imaging of vascular permeability in optically clear zebrafish embryos using fluorescently labeled tracers^28,29^. Additionally, the widespread recent emergence of CRISPR/Cas9 genome editing techniques have enhanced the study genetic regulators of endothelial barrier function in genetically engineered zebrafish mutants.

In response to a variety of cytokines, growth factors, and hormones, STAT proteins are activated via phosphorylation, dimerize, translocate to the nucleus, and bind to specific target gene promoters to regulate cellular processes, such as proliferation, differentiation, migration, and survival^21^. Reports have demonstrated VEGF rapidly induces STAT3 tyrosine phosphorylation and nuclear translocation in microvascular endothelial cells^22-24,56^. A positive feedback loop exists as VEGF-induced STAT3 has been shown to be a direct transcriptional activator of the VEGF promoter^57,58^. Functionally, phosphorylation of STAT3 by VEGF/VEGFR-2 signaling is required for endothelial cell migration^24^. Here, we show the integral role of STAT3 in the molecular regulation of vascular permeability using VEGF-inducible zebrafish crossed to CRISPR/Cas9-generated STAT3 mutants. Genomic ablation of STAT3 substantially reduces VEGF-mediated extravasation of fluorescent dextran from intersegmental vessels within the trunk region of larval zebrafish. Correspondingly, genetic knockout of STAT3 in the endothelium of mice increases vascular barrier integrity. Pharmacological inhibition of STAT3 using PYR, AQ, and C188-9 reduces VEGF-mediated vascular permeability in wildtype mice and prevents tight junction disorganization typically caused by VEGF stimulation of cultured human endothelial cells. VEGF/VEGFR-2 signaling results in JAK2-mediated activation of STAT3, which enables STAT3 to translocate to the nucleus and transcriptionally regulate genes involved in vascular barrier integrity, including intercellular adhesion molecule 1 (ICAM-1). Here, we identify ICAM-1 as a target of STAT3 transcriptional regulation. ICAM-1, a cell surface glycoprotein, has been shown to mediate VEGF-induced vascular permeability and leukostatsis^59^. IFNγ stimulation induces expression of both membrane-associated and soluble forms of ICAM-1, the latter of which binds to lymphocyte function associated antigen-1 to regulate immunomodulation^60^. ICAM1 is upregulated during inflammation stimulated by NF-κB or TNFα. In a rat lung injury model, ICAM-1 was suppressed via inhibition of TNFα- and IL-6-induced JAK2/STAT3 activation through dexamethasone treatment^61^.

While genomic STAT3 deficiency in mice results in embryonic lethality, the endothelium tissue-specific STAT3 knockout mice we report here are healthy and fertile, which coincides with previous findings^62^. Correspondingly, we observe normal vascular development in CRISPR/Cas9-generated zebrafish with homozygous genomic STAT3 deficiency, visualized by fluorescent microangiography. *In vitro* studies suggest that a dominant-negative form of STAT3 suppresses human dermal microvascular endothelial cell tube formation on Matrigel and collagen^24^. However, endothelial cells isolated from endothelium-specific STAT3 knockout mice and cultured *ex vivo* initiate normal tube formation^62^. While endothelial cell-specific STAT3 knockout mice undergo physiologically normal developmental angiogenesis, these mice exhibit defects in tissue repair and decreased recovery from vascular injuries, including myocardial infarction, cerebral ischemia, and ischemia-reperfusion injuries^63-65^. These observations highlight the need to understand the temporal dynamics through which STAT3 regulates pathological vascular permeability and edema as well as restorative angiogenesis to repair damaged tissue.

The functional role of STAT3 in VEGF-induced permeability has not been directly investigated. Prior studies suggest other permeability inducers, such as IL-6, IgG, IgE, histamine, lipopolysaccharides, and eotaxin, mediate vascular permeability through STAT3 signaling^25,66-72^. For example, an *in vitro* study using human endothelial cells recently demonstrated that IL-6 promotes sustained loss of endothelial barrier function via STAT3 signaling^66^, and IL-6-induced STAT3 activation has been shown to induce vascular permeability in ovarian endothelial cells^71^. Furthermore, IL-6-induced retinal endothelial permeability was found to be dependent upon STAT3 activation in mouse retina^72^. Dominant negative STAT3 has been shown to reduce IL-6-induced vascular permeability associated with malignant pleural effusion in lung adenocarcinoma^25^ and decrease IgG-mediated vascular permeability during acute lung injury^70^. Patients and mice harboring STAT3 mutations, which cause autosomal dominant hyper-IgE syndrome, have been shown to be partially protected from anaphylaxis. HUVEC derived from patients with AD-HIES or treated with a STAT3 inhibitor exhibit decreased histamine- and IgG-E-mediated leakage^67^. Inhibition of STAT3 phosphorylation decreases LPS-induced myocardial vascular permeability in a murine model^69^. Eotaxin stimulates STAT3 phosphorylation and disrupts human endothelial barrier integrity^68^. Taken together with the important role of STAT3 in modulating VEGF-mediated vascular permeability in vertebrate models presented here, studies collectively suggest STAT3 is a central regulator of vascular barrier function.

Considerable effort has been devoted to the development of STAT3 inhibitors as therapeutic agents. In addition to the important role of STAT3 in the pathogenesis of vascular disease, ischemia, and other tissue injuries, STAT3 is aberrantly overexpressed in many human tumor types and correlates with poor cancer prognoses. Development of STAT3 peptide inhibitors designed to target the p-Y-peptide binding pocket within the SH2 domain stemmed from elucidating the STAT3β homodimer structure^73-76^. However, the clinical applicability of these STAT3 peptide inhibitors has been hindered by their lack of stability and inability to cross membranes. Non-peptidic small molecule inhibitors of STAT3 exhibited promising in vivo activity in preclinical studies, but most of these STAT3 inhibitors failed to progress to clinical trials because they required medium-to-high micromolar concentrations to achieve sufficient activity and necessitated additional optimization in order to be systemically administered to human subjects^77^. While developing and translating STAT3 inhibitors to the clinic has proven difficult because STAT3 is a transcription factor without intrinsic enzymatic activity^78^, several compounds that inhibit either the function or expression of STAT3 are currently in clinical trials^79^, including a decoy oligonucleotide that competitively inhibits STAT3 interactions with its target gene promoter elements^80^ and an antisense oligonucleotide inhibitor of STAT3 expression, AZD9150^81^.

Pyrimethamine (PYR; Daraprim®) is a clinically available, FDA-approved agent that directly inhibits STAT3-dependent transcription. This anti-microbial drug was discovered as a new STAT3 inhibitor based on its ability to oppose the gene expression signature of STAT3^40,42^. PYR inhibits STAT3 phosphorylation and transcriptional activity at micromolar concentrations known to be routinely achieved in humans without toxicity^40^. In the current study, PYR suppresses STAT3 activation in endothelial cells as we observe decreased phosphorylation of STAT3 Y705 by immunofluorescence and immunoblotting. PYR treatment prevents VEGF-induced ZO-1 disorganization, which suggests that STAT3 inhibition improves tight junction instability in endothelium. We demonstrate that PYR administered to zebrafish and mice substantially reduces VEGF-induced vascular permeability. Atovaquone (AQ; Mepron®), another FDA-approved anti-microbial agent with a strong human safety profile, has been shown to rapidly and specifically downregulate cell-surface expression of glycoprotein 130, which is required for STAT3 activation in multiple contexts^41^. Like PYR, we show that AQ stabilizes endothelial tight junctions in the presence of VEGF stimulation. We also show that STAT3 peptide inhibitor, C188-9, reduces VEGF-induced vascular permeability in mouse models and cultured human endothelial cells. Our collective studies using three different STAT3 inhibitors suggest that suppression of STAT3 activity protects the endothelial barrier from VEGF-mediated vascular permeability. Future studies testing compounds that inhibit STAT3 activity, particularly PYR and AQ given their clinical accessibility, in models of human diseases involving pathological vascular permeability are warranted.

## Acknowledgements

This work was supported by a Grant-in-Aid of Research, Artistry and Scholarship Program Award #380634 from the Office of Vice President of Research at the University of Minnesota, Institutional Research Grant #129819-IRG-16-189-58-IRG81 from the American Cancer Society, and The Hormel Foundation (to L.H.H.) as well as the Fifth District Eagles Cancer Telethon Postdoctoral Fellowship Award (to S.K.A.). We thank Dr. Anna C. Sundborger-Lunna, Dr. Veer Bhatt, Dr. George Aslanidi, and Dr. Karina Kortova at The Hormel Institute for technically supporting the purification the human STAT3 protein and sharing associated reagents. We are grateful to Christina E. Hernandez, Abbygail M. Coyle, and Erin N. Dankert for their contributions to this work as Summer Undergraduate Research Experience interns. We thank Dr. Jim Hu from the Hospital for Sick Children in Toronto, Ontario, Canada for generously sharing the pGL3-ICAM1 reporter vector. We thank The Hormel Institute and its staff for administrative, shared equipment, animal facility, and institutional support.

## Author contributions

L.W. and M.A. performed vascular permeability studies in transgenic animals. L.W. completed vascular permeability studies in wildtype animals treated with pharmacological inhibitors. L.W. and M.A. conducted immunofluorescence and immunoprecipitation experiments. L.W., M.A., and Z.Z. completed immunoblotting studies. L.W. and S.K.A. performed in vitro kinase and luciferase transcriptional reporter assays. W.P. and S.M.B. generated and characterized STAT3 knockout zebrafish using CRISPR/Cas9 technology. D.A.F. provided technical and scientific support as well as shared compounds. L.W., M.A., S.K.A. and L.H.H. performed experimental troubleshooting, reviewed relevant scientific literature, and critically analyzed data. L.W. and L.H.H. prepared most figures and wrote the manuscript. L.H.H. conceived the original aims of this study, led the project, and acquired funding to complete the reported research. All authors approved the final version of this manuscript.

## SUPPLEMENTAL METHODS

### Cell Culture

Human endothelial cells were cultured in plates that had been pretreated for 30 minutes with Collagen I (Corning, Catalog No. 354231). HUVEC were maintained in EBM Endothelial Cell Growth Basal Medium (Catalog No. CC-3121, Lonza) supplemented with EGM Endothelial Cell Growth Medium SingleQuots (Catalog No. CC-4143, Lonza). HPAEC were cultured in EBM-2 Basal Medium (Catalog No. CC-3156, Lonza) supplemented with EGM SingleQuots (Catalog No. CC-4176, Lonza). HMVEC-L were cultured in EBM-2 Basal Medium (Catalog No. CC-3156, Lonza) supplemented with EGM SingleQuots (Catalog No. CC-4147, Lonza). For in vitro studies, HUVEC, HPAEC or HMVEC-L grown in culture to approximately 80% confluence were serum starved for 16 hours and subsequently stimulated with human recombinant VEGF-165 protein (25 ng/ml; R&D Systems; MNPHARM) for indicated durations. When applicable, cells were treated with inhibitors or control vehicle for various indicated periods of time following serum starvation and preceding VEGF-165 protein stimulation. For an indicated subset of experiments, HUVEC were maintained in a 1:1 mixture of astrocyte conditioned medium (ScienCell Research Laboratories, Catalog No. 1811) and HUVEC standard medium.

### Antibodies

Immunoblotting was performed using antibodies were purchased from Cell Signaling Technology (CST) to detect ZO-1 (Catalog No.13663; dilution 1:1000), phosphorylated VEGFR-2 (Tyr1175, Catalog No. 2478; dilution 1:1000), phosphorylated STAT3 (Tyr705, Catalog No. 9145; dilution 1:500), phosphorylated JAK2 (Tyr 1007/1008, Catalog No. 3771; dilution 1:500), phosphorylated JAK1 (Tyr 1034/1035, Catalog No. 3331S; dilution 1:500), phosphorylated TYK2 (Tyr1054/1055, Catalog No. 9321S; dilution 1:500), VEGFR-2 (Catalog No. 2479; dilution 1:1000), STAT3 (Catalog No. 12640; dilution 1:1000), JAK2 (Catalog No. 3230; dilution 1:500), JAK1 (Catalog No. 3332; dilution 1:500) and TYK2 (Catalog No. 14193; dilution 1:500). Additional antibodies were obtained from Santa Cruz Biotechnology to detect phosphorylated STAT3 (Tyr705, Catalog No. sc-8059; dilution 1:500), ICAM1 (Catalog No. sc-18853; dilution 1:500), and tubulin (Catalog No. sc-5286; dilution 1:500). The monoclonal antibody for detecting GST (Catalog No. MA4-004; dilution 1:1000) was bought from Thermo Fisher Scientific. The monoclonal antibody against phospho-STAT3 (Tyr708, Catalog No. D128-3) in zebrafish was obtained from MBL International Corporation. Horseradish peroxidase-conjugated anti-rabbit (Catalog No. 7074; dilution 1:5000) and anti-mouse (Catalog No. 7076; dilution 1:5000) secondary antibodies (1 μg/μl) were purchased from Cell Signaling Technology.

Immunofluorescence was performed using antibodies against ZO-1 (CST, Catalog No.13663; dilution 1:200), p-STAT3 (Santa Cruz Biotechnology; Tyr705, Catalog No. sc-8059; dilution 1:50) or STAT3 (CST, Catalog No. 12640; dilution 1:100). The cells were then washed with PBS and incubated with CF™ 488A goat anti-rabbit secondary antibody (Sigma-Aldrich; Catalog No. SAB4600389; dilution 1:1000) or CF™ 594 goat anti-mouse secondary antibody (VWR; Catalog No. 20110; dilution 1:1000).

### Immunoprecipitation

HUVEC were lysed in RIPA buffer (Millipore) supplemented with protease inhibitor cocktail (Roche) and phosphatase inhibitor cocktail set V (Sigma). After quantifying protein using the Quick Start Bradford protein assay (Bio-Rad), 500 µg protein lysate was loaded to the supplied spin column and immunoprecipitation was achieved following the manufacturer’s protocol (Catch and Release Immunoprecipitation Kit; Catalog No.: 17-500; Millipore).

### Genotyping

#### Mice

Genotyping primer pairs for genotyping the Stat3^tm1Xyfu^/J mice are forward primer 5’-TTGACCTGTGCTCCTACAAAAA-3’ and reverse primer 5’-CCCTAGATTAGGCCAGCACA-3’. The genotyping primer pairs for Tg(Tek-cre)1Ywa/J mice are forward primer 5’-CGCATAACCAGTGAAACAGCATTGC-3’ and reverse primer 5’-CCCTGTGCTCAGACAGAAATGAGA-3’.

#### Zebrafish

Following imaging of vascular permeability, each individual zebrafish was euthanized and genomic DNA was extracted using standard techniques for subsequent STAT3 genotyping using the following pair of primers: 5’-GGTCTTCCACAACCTGCTG-3’ and 5’-TAGACGCTGCTCTTCCCAC-3’.

### Compounds

Pyrimethamine (PYR; 75 mg/kg; Sigma Aldrich; Catalog No. 46706) or control vehicle was delivered to mice by intraperitoneal injection daily for 15 days. PYR was dissolved in dimethyl sulfoxide (DMSO) to a concentration of 167 mg/ml and then diluted in PBS before injection into mice. For zebrafish studies, 10µM PYR or vehicle (DMSO) was added to zebrafish embryo water for 3 days prior to assessment of vascular permeability and immunoblotting studies. Atovaquone (AQ; Sigma Aldrich; Catalog No. A7986), C188-9 (Sigma Aldrich; Catalog No. 573128), and PYR were each dissolved in DMSO and used for in vitro studies at the indicated concentrations. For in vivo studies, C188-9 was dissolved in DMSO at 208mg/ml and diluted in 5% dextrose in water (D5W) before injection. Mice received C188-9 (100 mg/kg) or vehicle (DMSO in D5W) by intraperitoneal injection daily for 1 week. For in vivo studies, Tyrphostin AG 490 (40 mg/kg, Sigma Aldrich; Catalog No.T3434) was dissolved in DMSO at the concentration of 250 mg/ml and diluted in PBS before injection. Mice received AG490 or vehicle (DMSO in PBS) by intraperitoneal injection daily for 1 week.

### Glutathione S-transferase (GST) pull-down assays

The cDNA fragment encoding full-length human STAT3 was subcloned from pLEGFP-WT-STAT3^1^ into the mammalian expression vector pEBG^2^ using KpnI and NotI restriction enzyme sites and the following subcloning primer pairs: forward: 5’-GGGGTACCTCGAGCTCAAGCTTCAGGATGG-3’ and reverse: 5’-ATAAGAATGCGGCCGCTCACTTGTAGTCCATGGGGGAGGTA-3’. pLEGFP-WT-STAT3 was a gift from Dr. George Stark (Addgene plasmid # 71450; http://n2t.net/addgene:71450; RRID:Addgene_71450)^1^. pEBG was generously shared by Dr. David Baltimore (Addgene plasmid # 22227; http://n2t.net/addgene:22227; RRID:Addgene_22227)^2^. Freestyle 293F cells (Thermo Fisher Scientific) grown in FreeStyle™ 293 Expression Medium (Thermo Fisher Scientific; 12338018) in an incubator at 37°C with 8% CO2 were transfected with pEBG-STAT3 plasmid according to the instructions of the Freestyle 293 Expression System (Thermo Fisher Scientific) and then harvested by centrifugation at 1500 × g for 10 minutes. Cell pellets were suspended in PBS containing protease inhibitor cocktail (Roche) and lysed on ice using 15 s pulses of sonication repeated 7 times with a Sonic Dismembrator, Model 100 (Fisher Scientific). Lysates in 1% Triton, 5 mM DTT were centrifuged at 12000g for 10 minutes at 4°C, and Glutathione Sepharose 4B (GE Healthcare) was used to bind the GST fusion proteins from the supernatant. GST protein-coated beads were incubated with pre-cleared HUVEC cell lysates at 4°C overnight. The bead-protein complexes were then washed 5 times with pre-chilled PBS, and the proteins were eluted using 10mM Glutathione elution buffer at room temperature. The proteins were boiled for 5 minutes in the Laemmli sample buffer and then analyzed by immunoblotting.

### Purification of human STAT3 protein

STAT3 cDNA (human α isoform, residues 1-770) was subcloned from pLEGFP-WT-STAT3^1^ into the RGS-6xHis-pcDNA3.1 plasmid^3^, which was a gift from Dr. Adam Antebi (Addgene plasmid # 52534; http://n2t.net/addgene:52534; RRID: Addgene_52534)^3^. The 6xHis-STAT3α fragment was cloned into the pFastBac™Dial vector, generously provided by Dr. George Aslanidi at the University of Minnesota, The Hormel Institute. The plasmid was first transformed into *E. coli* DH10 MultiBac. Single colonies were inoculated into 2 ml antibiotic LB broth containing 50 µg/ml Kanamycin, 7 µg/ml Gentamycin, and 10 µg/ml Tetracycline and grown at 37°C overnight. After isolating recombinant Bacmid DNA, we transfected Sf9 insect cells, grew cells in Gibco™ Sf-900™ II SFM medium (Gibco, Catalog NO.10902088) in a 27°C incubator, harvested P0 baculovirus stock, and amplified P1 and P2 baculovirus in a 27°C, 90 rpm shaker. Sf9 cells were infected with P2 baculovirus (MOI=10) and cells were harvested by centrifugation at 1500 rpm for 10 minutes after incubation for 3 days. The pellet was resuspended in binding buffer [50 mM Tris-HCl (pH 8.0), 500 mM NaCl, 2 mM MgCl2, 10% glycerol, 1mM Tris (2-Carboxyethyl) Phosphine (T-CEP), 10 mM Imidazole] supplemented with complete protease inhibitors (Roche) and then lysed by 7 cycles of sonication (Fisher Scientific; Sonic Dismembrator, Model 100) each consisting of constant pulse for 15s on ice. The lysate was cleared by centrifugation at 30,000 × g for 30 minutes at 4°C. The supernatant was bound to high performance HisTrap column (GE Healthcare) using binding buffer [Sodium phosphate 20 mM, NaCl 500 mM, Imidazole 20 mM and T-CEP 0.5 mM] and eluted with an imidazole gradient (0 to 500 mM). The protein was then concentrated and loaded onto a Superdex 200 size exclusion column equilibrated in 50 mM sodium phosphate, 150 mM NaCl and 0.5 mM T-CEP. Peak fractions were analyzed by SDS-PAGE.

### Quantitation of zebrafish vascular permeability

ImageJ software was used to quantitate the extent of Texas Red-dextran in the extravascular space. Mean gray value was measured after converting the image type to 8-bit gray, setting scale to pixels, inverting the image and identifying 4 areas of extravascular space in the middle of the zebrafish trunk region using drawing/selection tools to avoid intersegmental vessels evident by green fluorescent signal originating from FITC-dextran. The background gray value minus the average gray value of the four regions is used for statistical analysis.

## SUPPLEMENTAL FIGURES

**Supplemental Figure 1:**
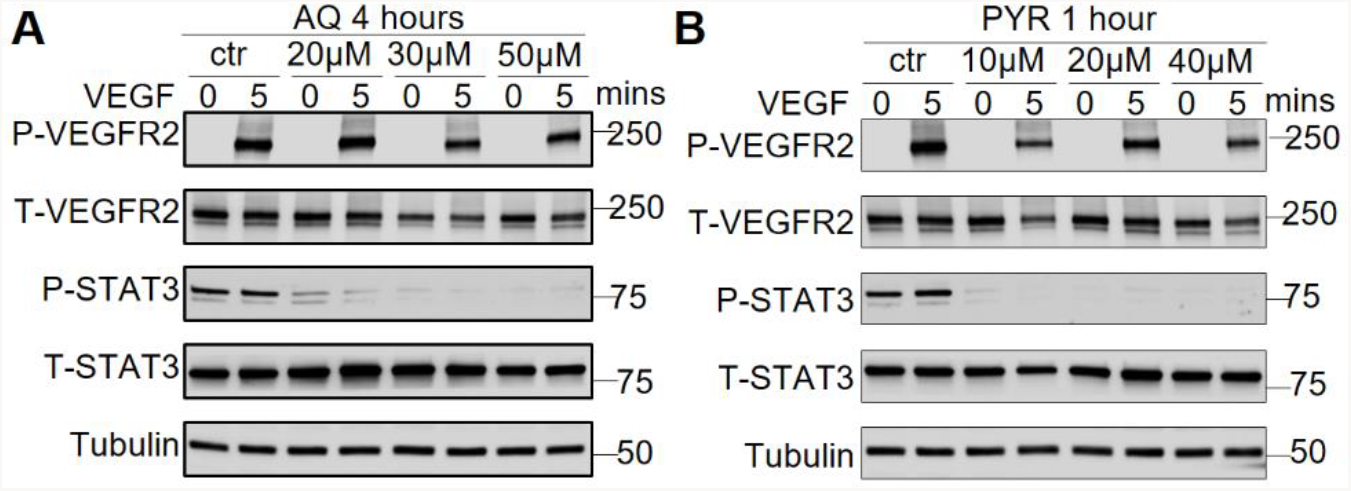
Assessment of STAT3 inhibition by atovaquone (AQ) and pyrimethamine (PYR) in VEGF-stimulated HUVEC. Serum-starved HUVEC were pretreated with 0, 20, 30, or 50 µM AQ for 4 hours (left) or 0, 10, 20, or 40 µM PYR for 1 hour (right) prior to human VEGF-165 protein (25 ng/ml) stimulation for 0 or 5 minutes. Cells were lysed and then immunoblotted with the indicated primary antibodies.

**Supplemental Figure 2:**
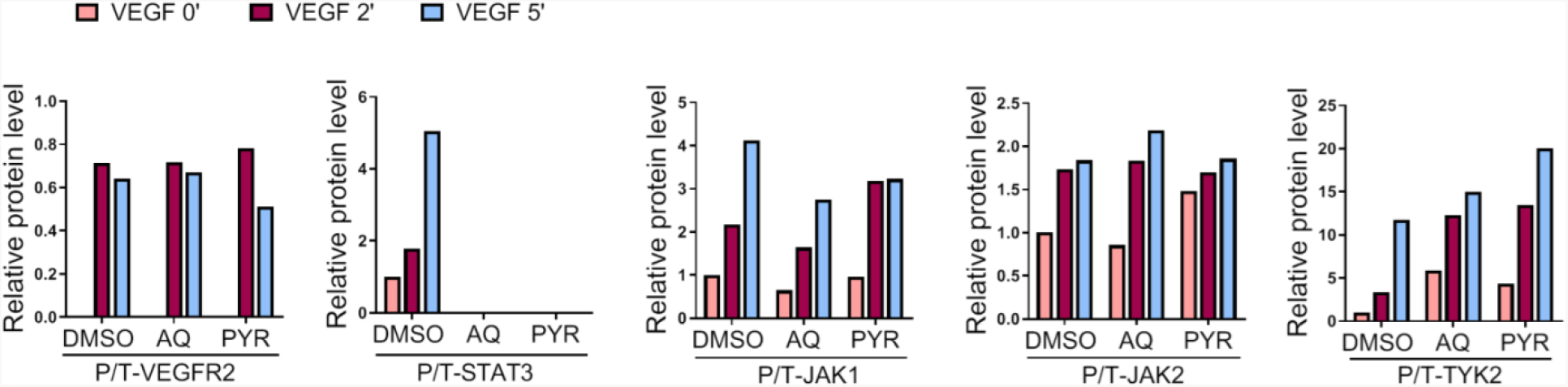
Densitometry analysis for immunoblotting of VEGF-stimulated HUVEC upon pretreatment with STAT3 inhibitors, atovaquone (AQ) and pyrimethamine (PYR). Densitometry was performed on the immunoblotting image depicted in Figure 4B by quantitating phosphorylated protein relative to total protein levels as indicated.

**Supplemental Figure 3:**
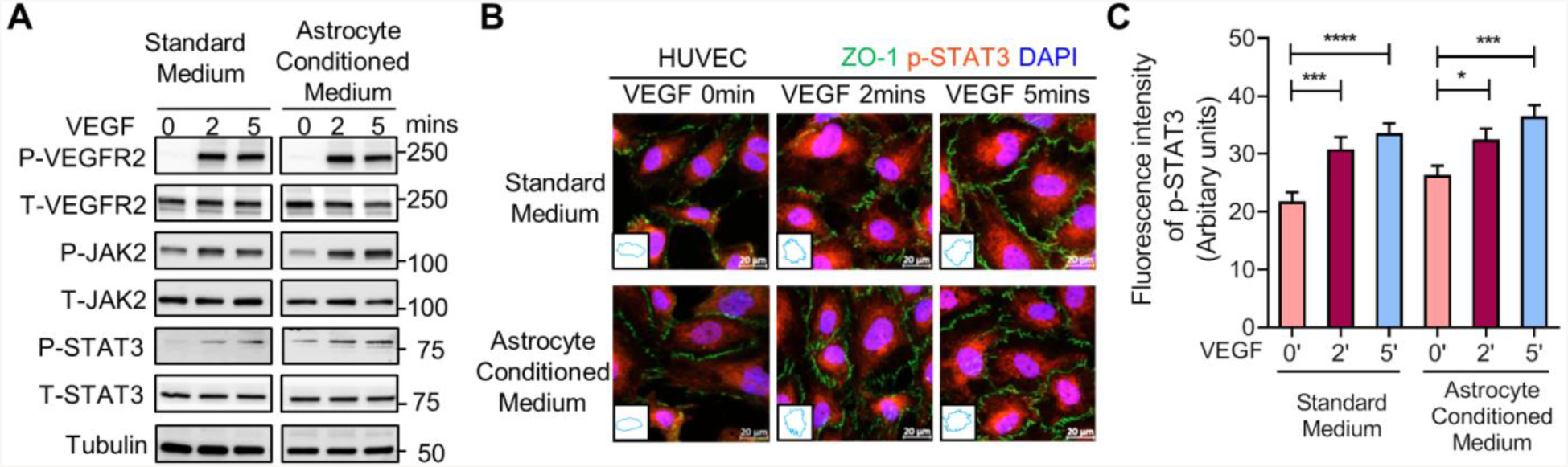
VEGF-induced STAT3 activation disrupts ZO-1 junctional stability in HUVEC. **A-B)** Serum-starved HUVEC were cultured in standard medium or astrocyte conditioned medium (mixed 1:1 with standard medium) and stimulated with VEGF (25 ng/ml). **A)** Cells were lysed and immunoblotted using indicated antibodies. **B)** IF was performed using p-STAT3 (Y705; red) and ZO-1 (green) antibodies. VEGF stimulation promotes disorganization of ZO-1 at endothelial cell junctions (i.e. jagged appearance). Nuclei stained with DAPI (blue). Insets: trace of ZO-1 staining. **C)** Quantification of the p-STAT3 staining intensity. *P<0.05, ***P<0.001, ****P<0.0001, one-way ANOVA.

**Supplemental Figure 4:**
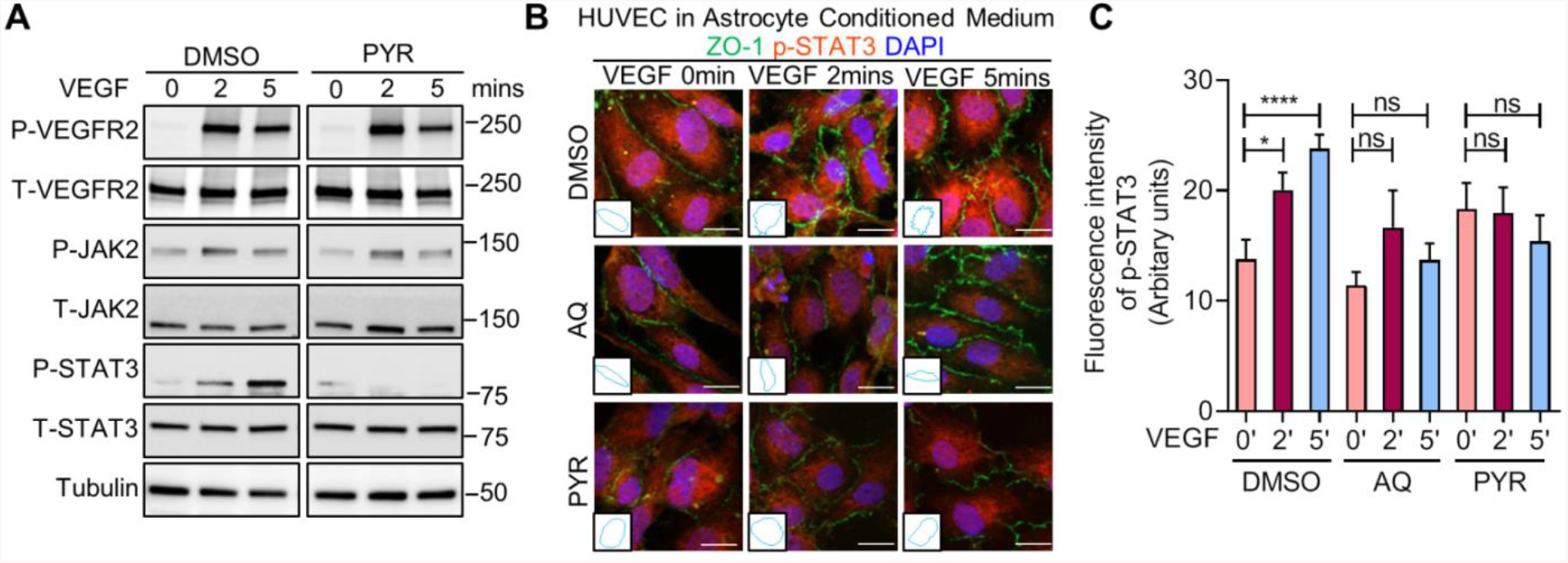
STAT3 inhibitors, atovaquone (AQ) and pyrimethamine (PYR), stabilize endothelial barrier integrity following VEGF stimulation in HUVEC cultured in astrocyte conditioned medium. **A)** Serum-starved HUVEC cultured in astrocyte conditioned medium (mixed 1:1 with standard medium) were pretreated with 20 µM PYR for 1 hour prior to VEGF (25 ng/ml) stimulation for 0, 2, or 5 minutes. **B)** Serum-starved HUVEC cultured in astrocyte conditioned medium (mixed 1:1 with standard medium) were pretreated with 30 µm AQ for 4 hours or 10 µm PYR for 1 hour prior to VEGF (25ng/ml) stimulation. VEGF stimulation promotes disorganization of ZO-1 (green) at endothelial cell junctions (i.e. jagged appearance). ZO-1 organization is maintained when HUVEC are pretreated with AQ or PYR (i.e. smooth appearance). Nuclei were stained with DAPI (blue). Insets: trace of ZO-1 staining on 1 cell. **C)** Quantification of the intensity of phosphorylated STAT3 protein at Y705. *P<0.05, ****P<0.0001, one-way ANOVA.

**Supplemental Figure 5:**
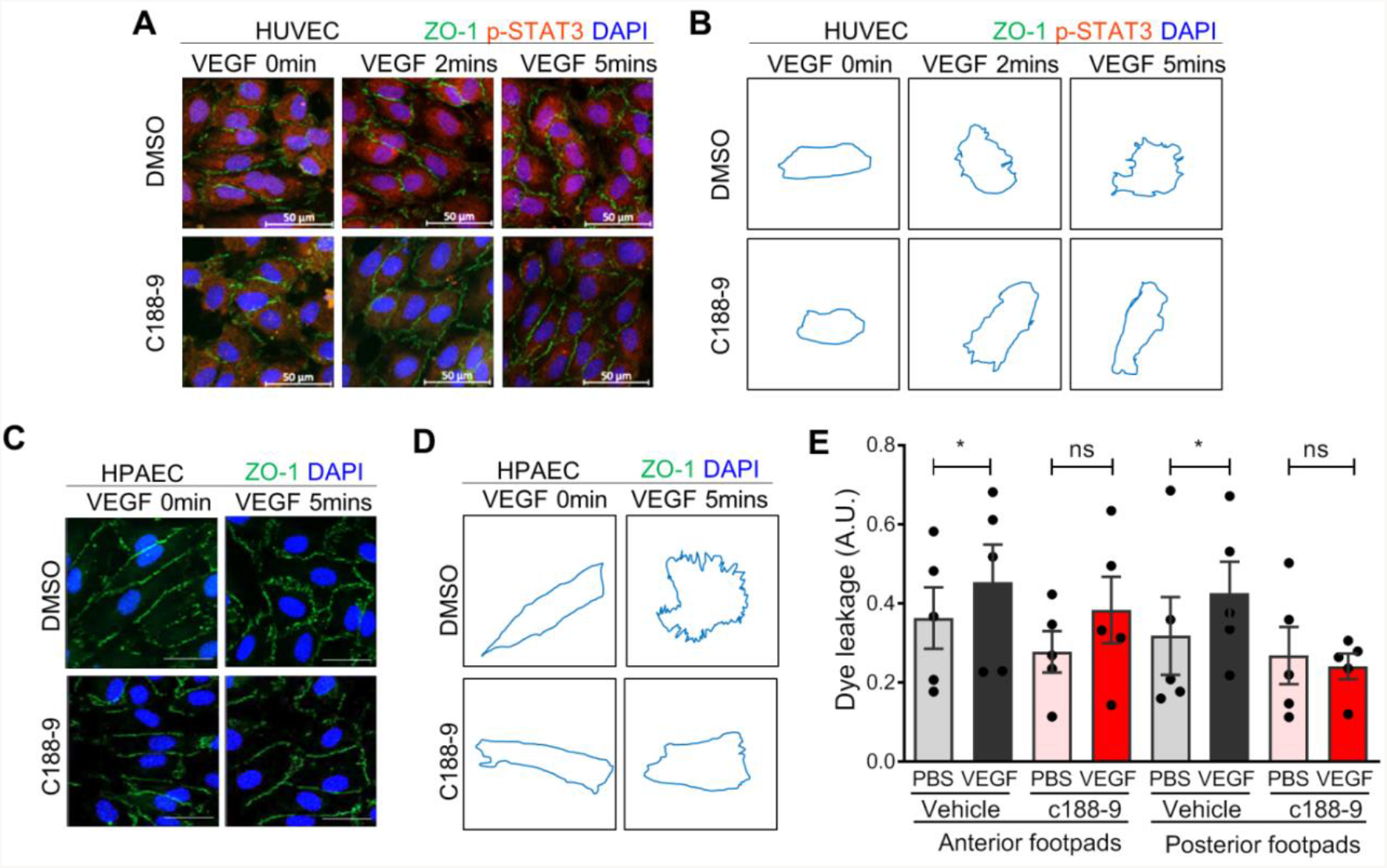
Pharmacological inhibition of STAT3 stabilizes endothelial barrier integrity following VEGF stimulation in HUVEC and mice. **A)** Serum-starved HUVECs were pretreated with 10 µM C188-9 for 5 minutes prior to human VEGF-165 protein (25ng/ml) stimulation for 0, 2 and 5 minutes. VEGF stimulation promotes disorganization of ZO-1 (green) at endothelial cell junctions. ZO-1 organization is maintained when HUVEC were pretreated with C188-9. p-STAT3 (Y705; red) was reduced upon treatment with C188-9. Nuclei were stained with DAPI (blue). **B)** Depiction of selected ZO-1 staining to help visualize its organization or disorganization upon VEGF treatment in the absence of STAT3 inhibitor, C188-9. **C)** Serum-starved HPAEC were pretreated with 10 µM C188-9 for 5 minutes prior to human VEGF-165 protein (25ng/ml) stimulation for 0 and 5 minutes. After stimulation with VEGF protein for 5 minutes, the structure of tight junction marked with ZO-1 was disrupted (i.e. jagged-like ZO-1 green staining). ZO-1 organization is maintained when HPAEC were pretreated with C188-9. Nuclei were stained with DAPI (blue). **D)** Depiction of selected ZO-1 staining. (E) Mice were administered C188-9 or vehicle prior to tail vein injection with 1% Evans blue and VEGF (2.5 µg/ml) or PBS vehicle being injected into the root of the footpad. Quantitation of Evans blue leakage in C57BL/6 wildtype mice. n=5 mice per group. Each mouse was injected with PBS on right anterior and posterior footpads and VEGF on left anterior and posterior footpads. Multiple biological replicates were performed and depicted findings are representative. *P<0.05, paired t-test.

**Supplemental Figure 6:**
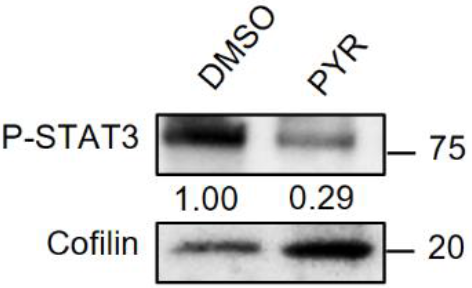
Pyrimethamine (PYR) inhibits STAT3 activity in zebrafish. Zebrafish were exposed to embryo water containing DMSO or 25 μM PYR for 3 days at starting 6 hours post-fertilization. Protein lysates were harvested from 3 days post-fertilization embryos by sonication in RIPA buffer after removing yolk sac and immunoblotting was performed using antibodies against zebrafish p-STAT3 (Y708) and cofilin.

